# Metabolic cost for isometric force scales nonlinearly and predicts how humans distribute forces across limbs

**DOI:** 10.1101/2023.12.24.573267

**Authors:** Sriram Sekaripuram Muralidhar, Nadja Marin, Colin Melick, Aya Alwan, Zhengcan Wang, Ross Baldwin, Sam Walcott, Manoj Srinivasan

**Author notes:** Co-first authors (Sriram Sekaripuram Muralidhar, Nadja Marin).

## Abstract

Muscles consume metabolic energy for force production and movement. A mathematical model of metabolic energy cost will be useful in predicting instantaneous costs during human exercise and in computing effort-minimizing movements via simulations. Previous in vivo data-derived models usually assumed either zero or linearly increasing cost with force, but a nonlinear relation could have significant metabolic or behavioural implications. Here, we show that metabolic cost scales nonlinearly with joint torque with an exponent of about 1.64, using calorimetric measurements of isometric squats. We then demonstrate that this metabolic nonlinearity is reflected in human behaviour: minimizing this nonlinear cost predicts how humans share forces between limbs in additional experiments involving arms and legs. This shows the utility of the nonlinear energy cost in predictive models and its generalizability across limbs. Finally, we show mathematical evidence that the same nonlinear metabolic objective may underlie force sharing at the muscle level.

---

Muscles consume metabolic energy to perform voluntary (walking, running, reaching, etc.) and involuntary movement (heart contraction, gastric motility, etc.). There is substantial evidence that humans move and act in a manner minimising the metabolic energy consumption for the task [1, 2, 3, 4], sometimes termed the energy cost. Hence, researchers interested in understanding human movement behaviour would benefit from methods for estimating the metabolic cost of human movement. But the most popular experimental technique to measure in vivo muscle energy cost, namely, indirect calorimetry, cannot measure instantaneous costs and requires the task to be repeated for 5-6 mins [5] or requires extrapolation from non-steady state data [6]. A mathematical model of the metabolic energy cost can be broadly useful in estimating energy costs of dynamic tasks, potentially in real-time, which cannot be repeated for extended periods and in reducing the time of cost estimation. Such models could enable novel studies of transient tasks [3] and faster development of human assistive devices like exoskeletons and prostheses, which are often evaluated based on their energy savings [7, 8]. Energy cost could also be useful either to predict human behaviour in a novel task or estimate the energy cost during simulations of normal, pathological and externally assisted human tasks. Here, our goal is to perform metabolic human subject experiments to obtain an energy cost model for isometric force production, and then test via further human experiments what human behaviour is predicted by minimizing this energy cost model.

We focus on developing an energy cost model for a constant force isometric task, that is, a task in which there is no movement but only force production with the muscle lengths being constant. We focus on isometric metabolic consumption because many slow and sub-maximal tasks may be dominated by the cost of isometric force, and the metabolic literature’s focus has been more on non-isometric tasks. For instance, previous models derived from in vivo experiments either predict zero cost for constant force isometric task (as their focus was non-isometric tasks with muscle length changes and mechanical work [9, 10, 11]) or usually assumed linear relations with isometric muscle force [12], or did not explicitly seek a nonlinear relation of energy cost with joint torque or force level [12, 13, 14] with one exception [15]. Some previous in vivo studies showed dependence of isometric metabolic cost with force rate and force exertion frequency [13, 16] but did not consider dependence with isometric force level. Previous metabolic models based on in vitro muscle experiments have not been compared with in vivo isometric force tasks [17, 18, 19, 20, 21]. Here, we perform new human subject experiments involving near-isometric squat experiments and fit a nonlinear relation with torque to the resulting data, showing that the nonlinear relation captures the data better than linear models (figure 1A). We specifically sought power law relations between metabolic cost and force, as there is precedence for using power law effort costs to solve the muscle force indeterminacy (sharing) problem [22], as well as power laws’ ability to potentially be applicable across scale and capture self-similarity [23].

**Figure 1:**
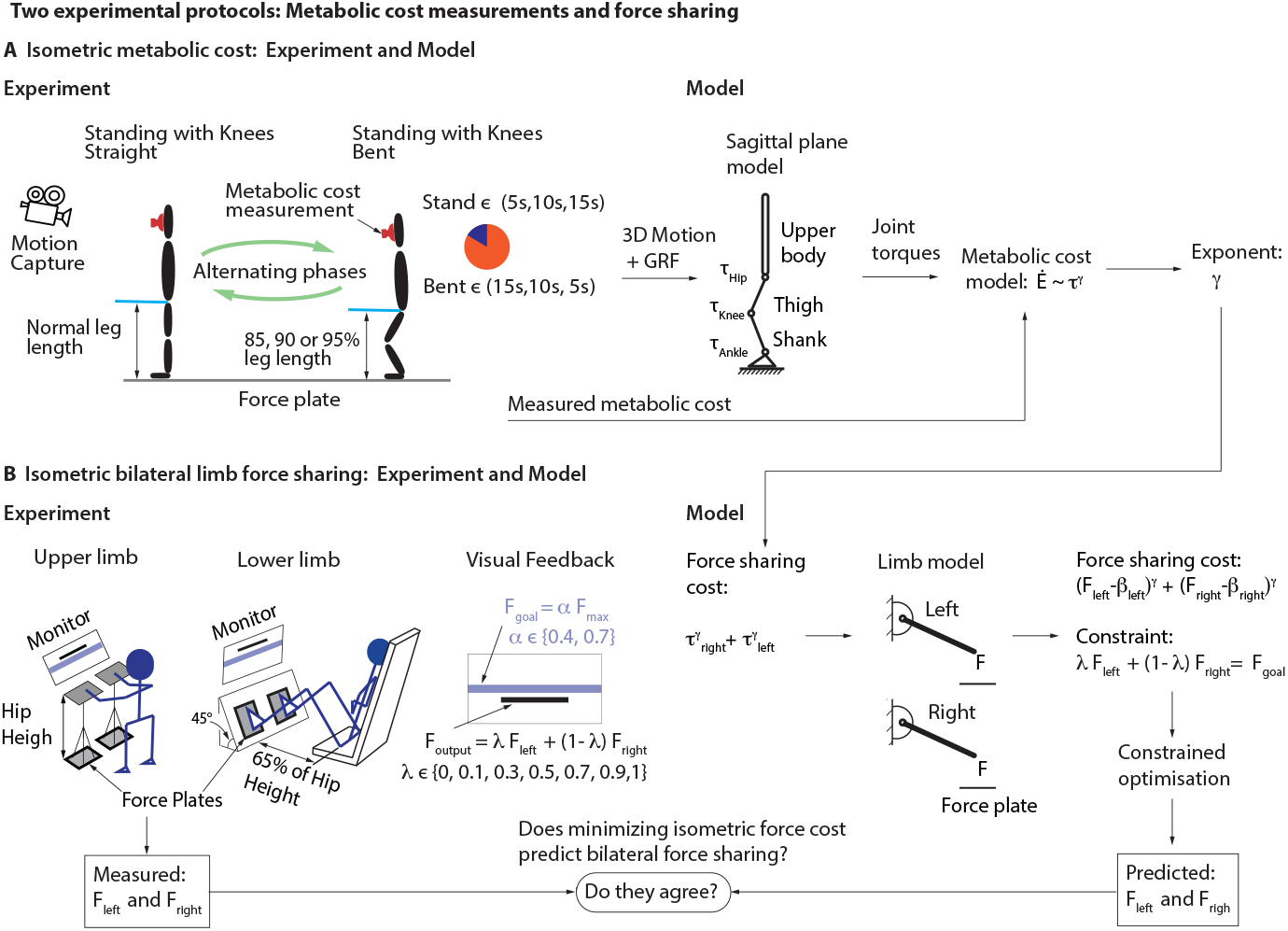
Two experimental protocols and two models: metabolic cost measurements and force sharing. A) Experiment: Human subjects alternate between quiet standing and squatting for 6 minutes using visual feedback. Squatting height (85, 90 or 95% leg length) and stand-squat durations ([5s,15s],[10s,10s] or [15s,5s]) are fixed for a trial and varied across trials. We estimated metabolic cost based on respiratory gases, 3D motion from optical motion capture, and ground reaction force from force plates. **Model:** Three rigid segment sagittal plane model of the human with rigid shank, thigh, and upper body’s geometric and inertia properties obtained from subject mass and height based on standard scaling [31]. Joint torques estimated from ground reaction force and 3D motion data. Metabolic cost is expressed as a function of joint torque and the coefficients are estimated by fitting the model to the experimental energy cost **B) Experiment:** Human participants apply forces with their limbs to track a fixed goal force with the help of visual feedback. Setup: For upper limbs, the participants stand straight and press vertically down on the platforms. Platform’s height is matched with the hip height of the participant. For lower limbs, the participants sit comfortably with their hands folded and press roughly perpendicular to the force plate. Visual feedback: The goal force is depicted by violet rectangle (*±* 10% of goal force) and the participant’s total force output by a black line. The goal force (F_goal_) and the left-right force contribution (*λ*) are fixed in a trial and varied across trials. **Model:** Constrained optmisation of the metabolic cost function to predict force sharing. metabolic cost function expressed as a power law function of joint torque with the exponent estimated from metabolic measurements. The joint torques were expressed in terms of measured external force using a single rigid segment sagittal plane model of the limb. The segment geometric and inertia properties are shown for illustration purposes as they are determined from constrained optimisation of cost function.

To test whether optimization of our new energy cost objective predicts human behavior in isometric tasks, we performed new human behavioral experiments in which humans had a choice regarding how they shared forces between different limbs: that is, in bilateral limb force sharing tasks (figure 1B). Previous experimental studies have shown patterns in limb force sharing for a given external task constraint [24, 25, 26, 27, 28] and here, we explain such patterns via metabolic optimization. A mathematical model of force sharing in healthy humans generalizable to both upper and lower limbs will be useful in understanding human behaviour in a novel situation, designing rehabilitation therapies for populations with neurological diseases and design of anthropomorphic robots [29, 30].

Finally, while our metabolic cost model is at the level of human joints, we hypothesize that the joint-level power law relation may have its origins in a corresponding muscle-level power law relation between muscle force and metabolic cost. To support this hypothesis, we provide a mathematical proof showing that if muscle-level force cost scales in a power-law way, the same exponent will be reflected in how the metabolic cost of multi-joint force-exertion tasks scales with the applied force.

## Metabolic cost of isometric torques scales nonlinearly with torque

We estimated the metabolic cost of isometric forces via indirect calorimetry experiments in which human subjects alternated between quiet standing for some time and standing with knees bent for some time, continuing the task for 6-7 minutes for a reliable metabolic measurement (figure 1A, Experiment; see *Materials and Methods*).

The prolonged squat period (standing with knees bent) necessitates use of near isometric muscle forces and requires substantial effort. Metabolic cost of alternating between standing and squatting increases with the increase in the relative duration of the squatting (figure 2A). We sought to model this increased metabolic cost as a function of the joint torques *τ*, specifically employing a power law relation of the form *τ* ^*γ*^, with the goal of determining whether the best fit exponent *γ* was systematically different from unity, thus evaluating if a nonlinear model was better than a linear model.

**Figure 2:**
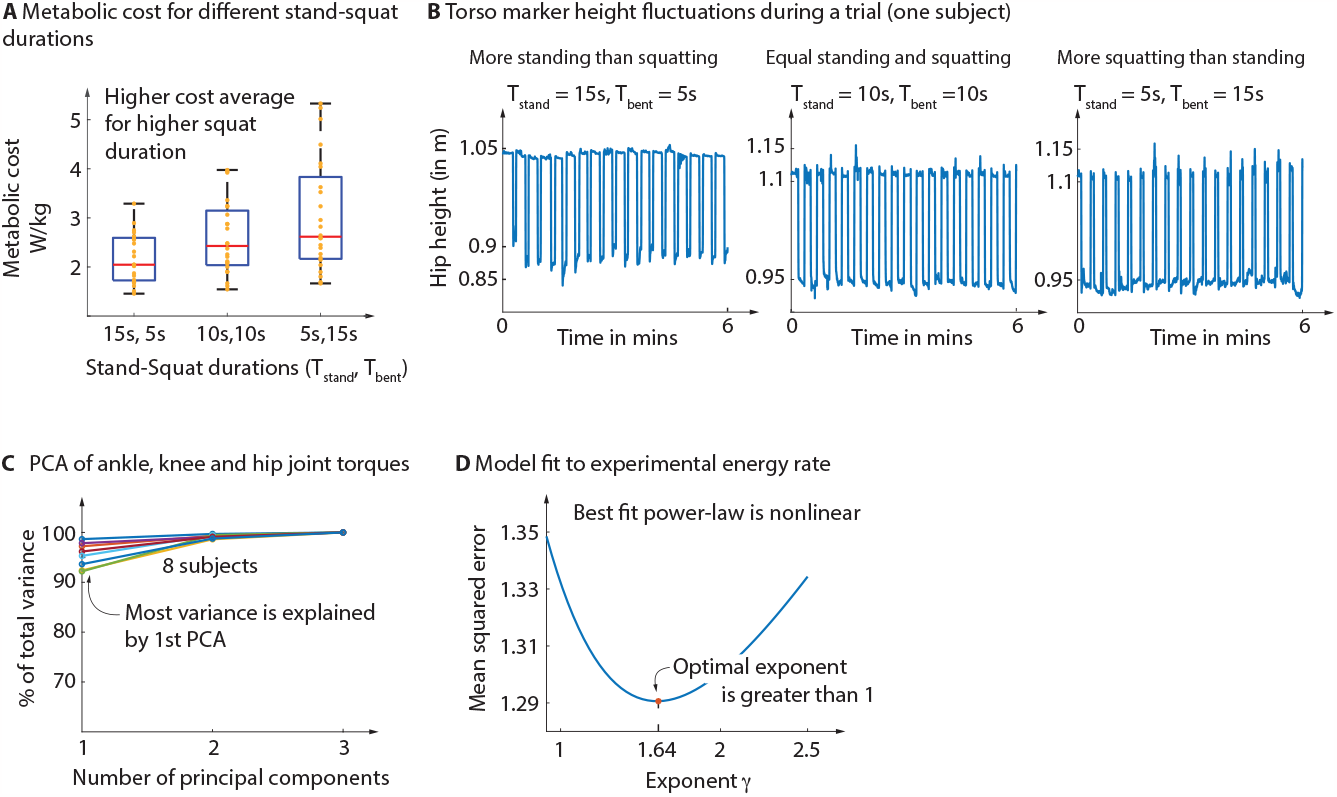
Metabolic cost of isometric tasks. **A)** Box plot of experimental metabolic cost (alternating between standing and squatting) for a given stand-squat duration with the data of 8 different subjects performing 3 different squat heights. More time spent in squat position results in more metabolic cost. **B)** Hip marker height data from the ground for a subject during 3 different trials. The movement resembles a square wave with different duty cycle with maxima denoting standing and minima denoting squatting. **C)** Principal component analysis of ankle, knee, and hip joint torques across all trials for each 8 subjects, percentage of variance explained by each principal component, showing the first component explains most of the data. **D)** The difference between model prediction and experimentally measured energy rate across all subjects and trials for different values of the model exponent (*γ*). The exponent with the least mean squared error is *γ* = 1.64.

Measured hip motion during the trials revealed that the subjects were reasonable but not perfect in alternating between the static postures throughout the 6 min trial (figure 2B). Nevertheless, this provided sufficient data for building the metabolic model. We performed inverse dynamics to estimate joint torques for the entire time duration. Principal component analysis of each subject’s ankle, knee and hip joint torque data of all trials showed that the first principal component was sufficient to explain over 90% of the data (figure 2C). So, the torques are mostly linearly related, and essentially proportional — because the torques are zero simultaneously, so the constant term in the linear relation among them must also be zero.

We fit the metabolic cost to the power-law energy cost model 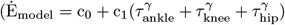), with a common coefficient (*c*_1_) for the three joint torques, because the torques being proportional across the data meant that, for the purposes of inferring the exponent (*γ*), the coefficients in the metabolic expression do not matter. The mean squared residual between model and experimental energy cost (averaged across all subjects and trials) revealed the optimum exponent *γ* to be 1.64 (figure 2D). Additionally, we obtained the best-fit *γ* for different relative coefficients (0.5 *<* b_0_, b_1_, b_2_ *<* 1.5) between the three joint torques in the following energy cost model: 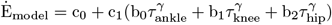. We found the optimum gamma values ranging between 1.5 and 1.8 which is consistent with the flatness of the MSE plot (figure 2D). This check gives us additional confidence in choosing equal coefficients for the different joint torques. Thus, the metabolic cost of isometric torques scales nonlinearly with torque, as *γ* = 1.64 is strictly better than linear dependence given by *γ* = 1.

### Bilateral limb force sharing is explained by metabolic minimization

To test whether the nonlinear metabolic cost model has behavioral implications, we performed bilateral limb force sharing experiments in which human subjects produced forces with their two hands or two legs, so as to achieve a fixed goal force (F_goal_) via a linear combination of the left and the right force F_output_ = *λ*F_left_ + (1 − *λ*)F_right_ (figure 1B). We considered two goal force levels, parameterized by *α*, the goal force normalized by the maximum comfortable force. See *Materials and Methods*. We hypothesized that humans selected the individual limb forces F_left_ and F_right_ in a manner that minimised energy cost. We expressed the force sharing cost as a power law function of joint torque 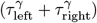 with the exponent (*γ*) derived from the previous metabolic energy cost measurements and performed constrained optimisation to predict the force sharing and compared with the experiments (figure 1B).

Subjects were good at tracking the goal force in both upper limb and lower limb experiments (figure 3A). In the lower limb force sharing experiments, for both goal force levels, subjects applied more force on the force plate which contributes more, quantified by *λ* — the extent of this asymmetry varied with *λ* and goal force level (figure 3D). Subjects in the upper limb force sharing experiment showed the same trends for the higher force level but this trend was less clear at the lower goal force, as the forces in the two limbs were close to being equal (figure 3C).

**Figure 3:**
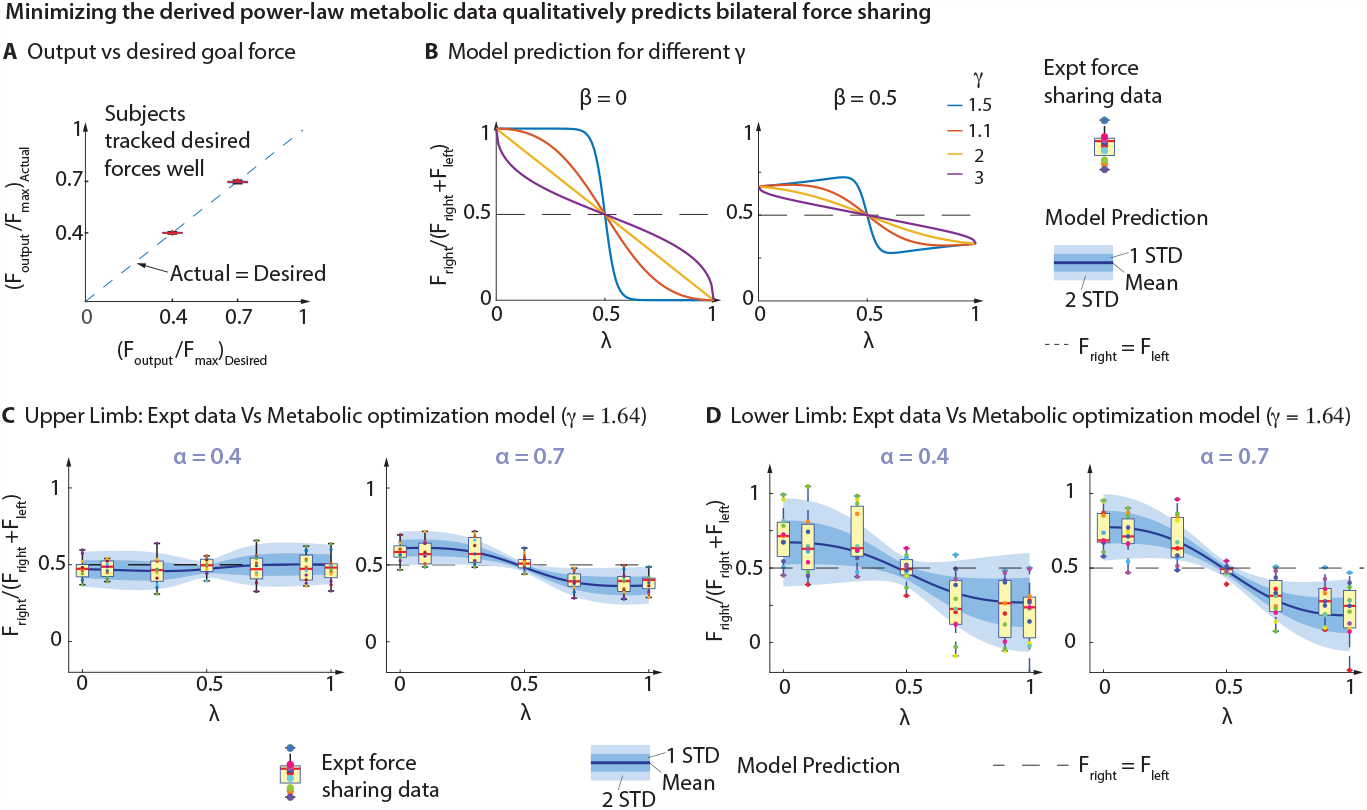
Bilateral force sharing: metabolic model prediction vs. experiment. Upper and lower limb bilateral force sharing. **A)** Boxplot of actual force tracked by the subjects averaged across second half of the trial in upper and lower limb experiments. **B)** Model prediction for different model parameters. **C)** Model with energy cost exponent 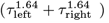 prediction overlaid on 12 human subject’s data for different left-right force plate contributions (*λ*) for lower (*α* = 0.4) and higher (*α* = 0.7) goal force levels. Lower limb bilateral force sharing. **D)** Model with energy cost exponent 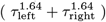 prediction overlaid on 12 human subject’s data for different left-right force plate contributions (*λ*) for lower (*α* = 0.4) and higher (*α* = 0.7) goal force levels.

Model predictions for substantially different exponents *γ* are qualitatively different (figure 3B), so this experiment allows us to test if the metabolically derived *γ* predicts the observed data. When force contribution on one side increases, the extent of asymmetry in force sharing increases and the magnitude is controlled by the parameter *β* (figure 3B), a measure of limb forces when the joint torques are zero, but the shape of the force sharing curve is mainly controlled by exponent *γ*. For instance, when *β* = 0, a nonlinear exponent of *γ* = 2 implies that force sharing changes are linear in *λ*, whereas when *γ* = 1 (linear metabolic rate), the predicted metabolic cost is indifferent to how the force is shared, so all force sharing strategies are optimal. When *γ* is just slightly greater than 1 (almost linear metabolic cost), the predicted force sharing is all or nothing (figure 3B), with one limb contributing the entire force depending on whether *λ* is greater or less than 0.5, but all other force sharing strategies are still close to energy optimal.

Using the exponent *γ* = 1.64 from the metabolic energy rate model and using optimal *β* values, we made predictions for upper and lower limb force sharing, which is in good agreement with the experimental data (figure 3C-D). Independently, by fitting the force sharing model to the data, we found the optimum *β* and *γ*. The optimum *γ* was 1.63 for upper limbs (almost the same as energy cost exponent 1.64) but 1.41 for the lower limbs, but as noted, *γ* = 1.64 did almost as well as 1.41 in explaining the data, suggesting a flat minimum (figure 3C-D). The model predictions are symmetric about *λ* = 0.5: that is, the predicted force sharing is the same for contribution of *λ* and 1-*λ*, except reversing left and right, and this symmetry is observed qualitatively in the behavioral data.

### Theorem: Metabolic power law scaling predicts a linear muscle force scaling strategy in a static task

We have framed the power law metabolic cost model thus far as being a function of joint torques. In this section, we generalize the metabolic cost to being a power law function of muscle force and show one behavioural implication of this generalization. Specifically, we consider the task of producing an external force of different magnitudes but fixed direction using a limb with multiple muscles and multiple joints, when the limb is at rest. We show that minimizing the power law metabolic cost results in a ‘linear scaling strategy’ for muscle forces (as seen in some finger force experiments [32]): that is, once the optimal solution is determined for one external force magnitude, the optimal solution for any other external force magnitude is simply scaling all the muscle force magnitudes by the same scalar factor. To be specific, consider a task in which a multi-joint (open-chain) limb with *m* muscles (figure 4A) is at rest and needs to apply a 3D external force **F**_ext_, which is a scaled version *μ***F**_0_ of some nominal 3D external force **F**_0_. We show that if the optimal muscle force magnitudes for the nominal external force **F**_0_ are given by **F**_mus0_ ∈ ℝ^*m*^, then the optimal muscle force magnitudes for any other external force *μ***F**_0_ are given by *μ***F**_mus0_, assuming the external force is much larger than limb weight due to gravity.

**Figure 4:**
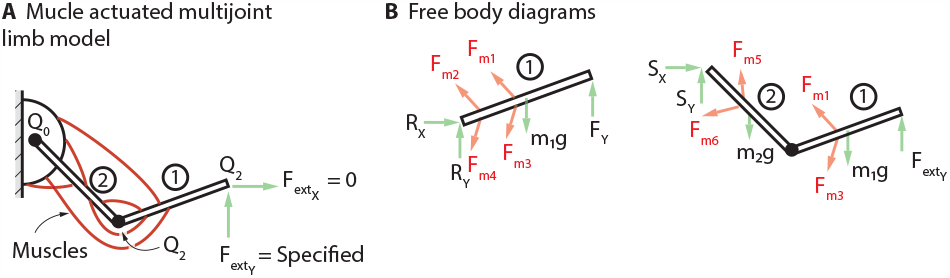
Multi-joint multi-muscle generalization: what muscle level metabolic scaling implies to joint level metabolic scaling.

The condition that the whole limb is in static equilibrium can be written as a linear equation in the list of *m* muscle force magnitudes **F**_mus_ ∈ ℝ^*m*^ as follows:

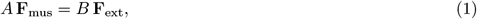

where *A* is a *p × m* matrix and *B* is a *p ×* 3 matrix, where the number of rows *p* equals the minimal number of scalar static equilibrium equations, generally equal to the number of degrees of freedom. In a static configuration, the muscle force directions are specified, so we only solve for the muscle force magnitudes **F**_mus_. The matrices *A* and *B* contain geometrical parameters such as instantaneous muscle moment arms and external force moment arms at the current limb configuration. Equation 1 ignores gravity, or equivalently, assumes that the external force to be produced is much larger than limb weight. For instance, for the two-segment limb of figure 4A, *p* = 2 with the equations obtainable by moment balance of the whole limb OAB about O and the segment AB about

A. We minimize the metabolic cost with a power law relation on muscle force magnitudes:

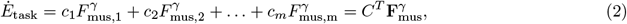

where the matrix *C* = [*c*_1_; *c*_2_; … *c*_*m*_] and 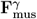 is defined as 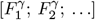. To solve the constrained optimization problem, we define the Lagrangian *L* as:

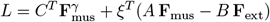

where *ξ* contains the Lagrange multipliers [*ξ*_1_; *ξ*_2_; … *ξ*_*p*_]. Differentiating this Lagrangian, that is, computing the gradient ∇*L* with respect to the unknown muscle force magnitudes **F**_mus_, and setting this derivative equal to zero gives:

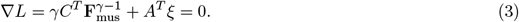

Equations 1 and 3 together provide *p* + *m* linear equations in the *p* + *m* unknowns in the *ξ* and **F**_mus_.

We now show that these equations 1 and 3 have solutions with an elegant structure that implies a simple scaling strategy for producing different external force levels. Assume that one needs to produce scaled versions of a fixed external force. That is, **F**_ext_ = *μ***F**_0_, where **F**_0_ ∈ ℝ^3^ is a fixed force and *μ* ∈ ℝ scales this force linearly to produce the external force **F**_ext_. We can rewrite the static equilibrium (equation 1) as:

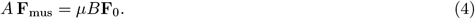

We now show that the solution is of the form:

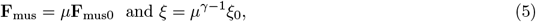

where **F**_mus_ is a fixed set of muscle force magnitudes. To do so, we substitute this solution form into equations 1 and 3, which gives the following two equations that are independent of *μ*:

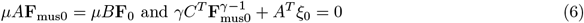

These equations allow us to solve for **F**_mus0_ and *ξ*_0_ without any dependence on *μ*, so how the muscle force magnitudes depend on *μ* is captured by equation 5. Thus, we predict that linearly scaling the external force is energy-optimally accomplished by linearly scaling the muscle force magnitudes. This linear scaling strategy is reminiscent of similar scaling observed in producing finger forces [32].

#### Corollary 1

**Muscle-level metabolic power law scaling predicts whole body level power law scaling with external force**. Substituting the muscle force solution (equation 5) into the metabolic cost expression (equation 2) shows that the total metabolic cost will scale like *μ*^*γ*^ :

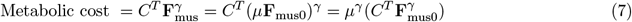

Given that the external force magnitude is |**F**_ext_| = *μ* |**F**_0_| is proportional to *μ*, the metabolic cost scales like external force raised to the power *γ*, when the external force is simply scaled.

#### Corollary 2

**Muscle-level metabolic power law scaling predicts whole body level power law scaling with external force**. At a given configuration, the joint torques are linear functions of muscle forces, given by:

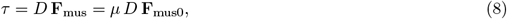

where *D* is a matrix of muscle moment arms. Thus, each of the joint torques is proportional to *μ* and scales linearly with the external force magnitude. So, when scaling the external force in a fixed configuration, the metabolic cost also scales like joint torque magnitude to the power *γ*. Recall that the theorem and the corollaries rely on fixing the configuration and having the external force be much higher than gravity, so gravity terms may be ignored. When gravity terms are substantial, the simple solution structure described above does not hold.

## Discussion

We found the metabolic cost of isometric force to scale non-linearly with joint torque (*τ* ^*γ*^), where the exponent *γ* was around 1.64. Our results of such nonlinearity agree qualitatively with previous in-vivo studies [33, 34, 35], which showed that oxygen consumption and the force are non-linearly related. Previous studies which suggested linear dependence of muscle metabolic cost with muscle force have either been for lower force range in in-vivo experiments [36, 37] or with isolated human heart or rat skeletal muscles in-vitro with external activation [38, 39, 40]. So, one possible source of the nonlinearity may be either the cost of activation and calcium pumping [41], and another could be the differential activation of motor units [42], with some muscle fibers producing more force than others and with fibers differing in properties (e.g., fast-twitch vs slow-twitch)[43, 44].

Minimizing the energy cost of force sharing with the cost scaling with torque *τ* as *τ* ^*γ*^ with exponent *γ* ≈ 1.64 predicted force sharing well in both upper limb and lower limb tasks. Our model is also in qualitative agreement with a previous bilateral upper limb force sharing experiment [27]. Terekhov et al [45] showed that the force sharing cost for five fingers force sharing can be explained by a quadratic function of force with a linear term, which is analogous to a power law with 1 *< γ <* 2 as suggested by our model. The same torque scaling for metabolic cost and force sharing cost suggests that metabolic energy optimality might be the reason for healthy human behaviour in bilateral limb force sharing. While we performed the metabolic experiments only in lower limbs, because the energy cost model satisfactorily predicted upper limb force sharing as well, we hypothesize that the metabolic cost model for upper limb joints may be similar.

The human body has more muscles than actuated degrees of freedom. This means that it is not possible to obtain muscle forces from inverse dynamics, only joint torques — sometimes called the redundancy problem. The classic force distribution problem in biomechanics involves computing how a given set of joint torques may be produced by appropriate muscle forces [22]. It is conventional to minimize an objective equal to the summed muscle stress raised to the power two, as done in OpenSim’s static optimization or computed muscle control [46]. We also showed analytically that the same exponent carries over to the muscle force sharing objective suggesting that the energy optimality might be the guiding factor for our brain to solve the muscle force indeterminacy problem for isometric contraction. This result may be more important than just for the force distribution problem – as it may be applicable more broadly to metabolic optimisation-based predictions.

We primarily focused on the muscle energetics scaling nonlinearity, estimating the exponent (*γ*), but we did not characterize the relative coefficients (*α*_i_) for each muscle in the force sharing objective. Future studies can focus on single joint experiments to estimate the coefficients while simultaneously measuring the energy cost to obtain direct comparisons. We do not have a microscopic explanation of the metabolic cost trends, and future work may consider explaining these via multiscale models that go from molecules, through sarcomeres, muscle fibers, recruitment, connective tissue, and whole body mechanics.

Limitations of our metabolic cost model include simplifications in both experiment and in the modeling. The model was derived from experiments where muscles are not perfectly isometric as there is a slight change in the length due to compliant nature of the tendon but this is the best we can achieve from in-vivo experiments. The metabolic rate analysis ignored the effect of muscle length or joint angle. Due to the approximate proportionality between joint torques, the analysis cannot determine the relative weight of the individual torques, which requires separate joint-specific experiments.

In conclusion, we have shown that energy cost scales non-linearly with joint torque and predicts bilateral limb force sharing and the same exponent is applicable to muscle force sharing. Such metabolic cost models, when generalized to each joint or muscle, may be used in whole body biomechanical simulations for predictions of movement behavior as well as potentially for real-time metabolic monitoring [47, 48].

## Materials and Methods

### Metabolic cost of an isometric task: Human metabolic experiments

We estimated the metabolic cost of isometric forces via experiments in which human subjects alternated between quiet standing for some time (T_stand_) and standing with knees bent for some time (T_bent_) with each trial lasting 6-7 minutes (figure 1A). The main idea is that when the subjects have their knees bent, while being still, the muscles are close to isometric. Metabolic cost 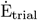 was estimated using Oxycon Mobile from measured volumetric rates of oxygen and carbon dioxide (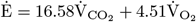 watt*/*kg [49]). Body movement was measured with a Vicon T20 motion capture system, and the ground reaction forces under each foot were measured with a Bertec instrumented treadmill. Eight subjects (all male; height= 1.81 *±* 0.05 m; mass= 77 *±* 11 kg; age= 24 *±* 6 years) participated with informed consent. Subjects were provided visual feedback to bend their knees to lower their hip to approximately 85-95% of their standing height. Three different T_stand_, T_bent_ combinations, namely {15 s, 5 s}, {10 s, 10 s}, {5 s, 15 s}, along with three prescribed knee-bend heights — approximately 85%, 90%, and 95% of normal leg length — gave nine squat trials per subject. In addition, we measured the cost of quiet standing (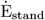) before the trials. The energy rate of squatting is estimated by subtracting the standing rate from the energy rate measured during the trial (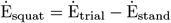). We exclude the cost of transition between standing to squatting in this above calculation for simplicity, as the transitions are a small fraction of the total task period.

### Metabolic cost of an isometric task: Mathematical model

We used a three-segment sagittal plane model of the human (figure 1A) with rigid shank, thigh, and upper body’s geometric and inertia properties obtained from subject mass and height based on standard scaling [31]. The foot remains largely motionless in the task, and thus does not contribute to the joint torques. We performed inverse dynamics to estimate the joint torques versus time for each trial, considering three joints: ankle, knee and hip. Subjects did not keep the knee bends exactly still, with some slow upward or downward motions that changed the knee angle, and the inverse-dynamics-based torques account for this movement. To each subject’s 9 trials, we sought to fit subject-specific metabolic rate model 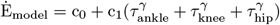 and estimate the coefficients c_0_, c_1_ for a range of fixed *γ* values by minimising the mean squared residual between the model and measured cost (Ė_squat_), averaged over the second half of the trial (3-6 mins). We fit this model with common coefficient (c_1_) for the three joint torques as the three joint torques were highly correlated within a trial (figure 2C). Once we obtain the best subject-specific model for each *γ*, we obtain the optimal exponent value *γ*_opt_ by minimising the squared residual between this model and measured cost, averaged across all subjects.

### Bilateral limb force sharing: Human behavioral experiments

We performed bilateral limb force sharing experiments in which human subjects produced forces with their two hands or two legs, so as to achieve a fixed goal force (F_goal_) via a linear combination of the left and the right force F_output_ = *λ*F_left_ + (1 − *λ*)F_right_ (figure 1B). This is a way of imposing force sharing in experiments which is a variant of previous studies [28, 50, 27]. We had upper and lower limb-specific setups with the individual limb forces (F_right_, F_left_) measured using force plates (Bertec, Vernier). Twelve subjects for upper limb (5M, 7F; height = 1.69 *±* 0.11m; mass = 68.83*±* 2.5 kg; age = 20.66*±*2.5 years; all right dominant) and twenty one subjects for lower limb (14M, 7F; height = 1.72 *±* 0.08m; mass = 68.1*±*10.22 kg; age = 23.6*±*4.82 years; mean*±*s.d; all right dominant) participated with informed consent and each performed 14 trials lasting 3 minutes each. We had seven different left-right force contributions (*λ* = *{*0, 0.1, 0.3, 0.5, 0.7, 0.9, 1*}*) and two different goal force levels which were a fraction (40 and 70%) of maximum limb force (F_max_). The maximum limb force was estimated from a 30 s trial where the subjects were instructed to apply a maximum force which could be held comfortably. We averaged force data from second half of the trial (15-30 s) to estimate the maximum left and right limb force, which was averaged to define F_max_. Subjects were told which side contributes more to the output for each trial but not the exact value of *λ*.

### Bilateral limb force sharing: Mathematical model

We hypothesized that humans selected the forces in a manner that minimised energy cost. We expressed the force sharing cost as a power law function of joint torque with the exponent (*γ*) derived from the previous energy cost measurements of isometric squat. We represented each limb using a single joint and a rigid segment in the sagittal plane (figure 1B). Hence, our force sharing metabolic cost is the sum of a power law function of the left and right limb joint torque: 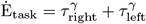. We expressed each joint torque in terms of measured limb force, segment weight, and limb proportions, as 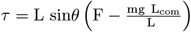, resulting in the following form of the energy cost (see also appendix 1):

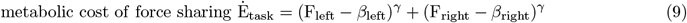

In this energy cost, *β*_left_, *β*_right_ was a subject specific parameter obtained by fitting to data and represents the forces exerted by the hands on the load cells due to gravity when the muscles are turned off entirely. The exponent *γ* was derived from the metabolic experiments. For each value of the parameter set, we performed optimisation with experimental constraint (*α*F_max_ = *λ*F_left_ + (1 − *λ*)F_right_) to predict the left and right limb force for all experimental trials. We did this separately for upper and lower limb experimental data to obtain a limb specific model. In addition to the predictions from the metabolically derived *γ*, we also performed additional optimizations to obtain the *γ* that best fit the experiments, while also allowing *β* to vary in a subject-specific way.

## Acknowledgments

This work was supported by NIH-R01GM135923 and NSF SCH grant 2014506.

## References

[1] M. Srinivasan, “Optimal speeds for walking and running, and walking on a moving walkway,” Chaos: An Interdisciplinary Journal of Nonlinear Science, vol. 19, p. 026112, June 2009. Publisher: American Institute of Physics.

[2] G. L. Brown, N. Seethapathi, and M. Srinivasan, “A unified energy-optimality criterion predicts human navigation paths and speeds,” Proceedings of the National Academy of Sciences, vol. 118, p. e2020327118, July 2021. Publisher: Proceedings of the National Academy of Sciences.

[3] N. Seethapathi and M. Srinivasan, “The metabolic cost of changing walking speeds is significant, implies lower optimal speeds for shorter distances, and increases daily energy estimates,” Biology Letters, vol. 11, p. 20150486, Sept. 2015. Publisher: Royal Society.

[4] L. L. Long and M. Srinivasan, “Walking, running, and resting under time, distance, and average speed constraints: optimality of walk–run–rest mixtures,” Journal of The Royal Society Interface, vol. 10, p. 20120980, Apr. 2013. Publisher: Royal Society.

[5] E. Ferrannini, “The theoretical bases of indirect calorimetry: A review,” Metabolism, vol. 37, pp. 287–301, Mar. 1988.

[6] J. C. Selinger and J. M. Donelan, “Estimating instantaneous energetic cost during non-steady-state gait,” Journal of Applied Physiology, vol. 117, pp. 1406–1415, Dec. 2014.

[7] S. K. Au, J. Weber, and H. Herr, “Powered Ankle–Foot Prosthesis Improves Walking Metabolic Economy,” IEEE Transactions on Robotics, vol. 25, pp. 51–66, Feb. 2009. Conference Name: IEEE Transactions on Robotics.

[8] S. H. Collins, M. B. Wiggin, and G. S. Sawicki, “Reducing the energy cost of human walking using an unpowered exoskeleton,” Nature, vol. 522, pp. 212–215, June 2015. Number: 7555 Publisher: Nature Publishing Group.

[9] J. Doke and A. D. Kuo, “Energetic cost of producing cyclic muscle force, rather than work, to swing the human leg,” Journal of Experimental Biology, vol. 210, pp. 2390–2398, July 2007.

[10] J. Doke, J. M. Donelan, and A. D. Kuo, “Mechanics and energetics of swinging the human leg,” The Journal of Experimental Biology, vol. 208, pp. 439–445, Feb. 2005.

[11] J. C. Dean and A. D. Kuo, “Energetic costs of producing muscle work and force in a cyclical human bouncing task,” Journal of Applied Physiology, vol. 110, no. 4, pp. 873–880, 2011. Publisher: American Physiological Society.

[12] D. Hawkins and P. Molé, “Modeling energy expenditure associated with isometric, concentric, and eccentric muscle action at the knee,” Annals of Biomedical Engineering, vol. 25, pp. 822–830, Sept. 1997.

[13] T. J. van der Zee and A. D. Kuo, “The high energetic cost of rapid force development in muscle,” Journal of Experimental Biology, vol. 224, p. jeb233965, May 2021.

[14] T. J. van der Zee, K. K. Lemaire, and A. J. van Soest, “The metabolic cost of in vivo constant muscle force production at zero net mechanical work,” Journal of Experimental Biology, vol. 222, no. 8, p. jeb199158, 2019.

[15] E. Kuroda, V. Klissouras, and J. H. Milsum, “Electrical and metabolic activities and fatigue in human isometric contraction.,” Journal of Applied Physiology, vol. 29, pp. 358–367, Sept. 1970.

[16] T. J. Van Der Zee, K. K. Lemaire, and A. J. “Knoek” Van Soest, “The metabolic cost of in vivo constant muscle force production at zero net mechanical work,” Journal of Experimental Biology, p. jeb.199158, Jan. 2019.

[17] L. J. Bhargava, M. G. Pandy, and F. C. Anderson, “A phenomenological model for estimating metabolic energy consumption in muscle contraction,” Journal of Biomechanics, vol. 37, pp. 81–88, Jan. 2004.

[18] H. Houdijk, M. F. Bobbert, and A. de Haan, “Evaluation of a Hill based muscle model for the energy cost and efficiency of muscular contraction,” Journal of Biomechanics, vol. 39, pp. 536–543, Jan. 2006.

[19] G. A. Lichtwark and A. M. Wilson, “A modified Hill muscle model that predicts muscle power output and efficiency during sinusoidal length changes,” Journal of Experimental Biology, vol. 208, no. 15, pp. 2831–2843, 2005. Publisher: Company of Biologists.

[20] A. E. Minetti and R. M. Alexander, “A Theory of Metabolic Costs for Bipedal Gaits,” Journal of Theoretical Biology, vol. 186, pp. 467–476, June 1997.

[21] B. R. Umberger, K. G. Gerritsen, and P. E. Martin, “A Model of Human Muscle Energy Expenditure,” Computer Methods in Biomechanics and Biomedical Engineering, vol. 6, pp. 99–111, May 2003.

[22] A. Erdemir, S. McLean, W. Herzog, and A. J. van den Bogert, “Model-based estimation of muscle forces exerted during movements,” Clinical Biomechanics, vol. 22, pp. 131–154, Feb. 2007.

[23] J. T. Kuikka, “Scaling laws in physiology: relationships between size, function, metabolism and life expectancy,” International Journal of Nonlinear Sciences and Numerical Simulation, vol. 4, no. 4, pp. 317–328, 2003.

[24] X. Hu and K. M. Newell, “Visual information gain and task asymmetry interact in bimanual force coordination and control,” Experimental Brain Research, vol. 212, pp. 497–504, Aug. 2011.

[25] Y. Jin, M. Kim, S. Oh, and B. Yoon, “Motor control strategies during bimanual isometric force control among healthy individuals,” Adaptive Behavior, vol. 27, pp. 127–136, Apr. 2019. Publisher: SAGE Publications Ltd STM.

[26] X. Hu and K. M. Newell, “Modeling constraints to redundancy in bimanual force coordination,” Journal of Neurophysiology, vol. 105, pp. 2169–2180, May 2011. Publisher: American Physiological Society.

[27] X. Hu and K. M. Newell, “Adaptation to bimanual asymmetric weights in isometric force coordination,” Neuroscience Letters, vol. 490, pp. 121–125, Feb. 2011.

[28] N. E. Skinner and A. D. Kuo, “Targeted limb rehabilitation using a reward bias,” Oct. 2018.

[29] T. Endo, T. Yoshikawa, and H. Kawasaki, “Multi-Fingered Bimanual Haptic Interface Robot with Three-Directional Force Display,” Advanced Robotics, vol. 25, pp. 1773–1791, Jan. 2011.

[30] T. C. Pataky, M. L. Latash, and V. M. Zatsiorsky, “Prehension synergies during nonvertical grasping, II: Modeling and optimization,” Biological cybernetics, vol. 91, pp. 231–242, Oct. 2004.

[31] P. de Leva, “Adjustments to Zatsiorsky-Seluyanov’s segment inertia parameters,” Journal of Biomechanics, vol. 29, pp. 1223–1230, Sept. 1996.

[32] F. J. Valero-Cuevas, “Predictive modulation of muscle coordination pattern magnitude scales fingertip force magnitude over the voluntary range,” Journal of Neurophysiology, vol. 83, no. 3, pp. 1469–1479, 2000.

[33] E. Kuroda, V. Klissouras, and J. H. Milsum, “Electrical and metabolic activities and fatigue in human isometric contraction.,” Journal of Applied Physiology, vol. 29, pp. 358–367, Sept. 1970.

[34] W. T. Josenhans, “Muscular factors.,” Canadian Medical Association Journal, vol. 96, pp. 761–764, Mar. 1967.

[35] P. Cerretelli, A. Veicsteinas, M. Fumagalli, and L. Dell’orto, “Energetics of isometric exercise in man,” Journal of Applied Physiology, vol. 41, pp. 136–141, Aug. 1976.

[36] D. H. Clarke, “Energy Cost of Isometric Exercise,” Research Quarterly. American Association for Health, Physical Education and Recreation, vol. 31, pp. 3–6, Mar. 1960.

[37] J. Royce, “Oxygen intake curves reflecting circulatory factors in static work,” Internationale Zeitschrift für angewandte Physiologie einschließlich Arbeitsphysiologie, vol. 19, pp. 222–228, July 1962.

[38] R. Paul and J. Peterson, “Relation between length, isometric force, and O2 consumption rate in vascular smooth muscle,” American Journal of Physiology-Legacy Content, vol. 228, pp. 915–922, Mar. 1975.

[39] E. Glück and R. J. Paul, “The aerobic metabolism of porcine carotid artery and its relationship to isometric force: Energy cost of isometric contraction,” Pflügers Archiv European Journal of Physiology, vol. 370, no. 1, pp. 9–18, 1977.

[40] D. A. Hood, J. Gorski, and R. L. Terjung, “Oxygen cost of twitch and tetanic isometric contractions of rat skeletal muscle,” American Journal of Physiology-Endocrinology and Metabolism, vol. 250, pp. E449–E456, Apr. 1986.

[41] D. W. Russ, M. A. Elliott, K. Vandenborne, G. A. Walter, and S. A. Binder-Macleod, “Metabolic costs of isometric force generation and maintenance of human skeletal muscle,” American Journal of Physiology-Endocrinology and Metabolism, vol. 282, pp. E448–E457, Feb. 2002.

[42] N. Fallentin, K. Jorgensen, and E. B. Simonsen, “Motor unit recruitment during prolonged isometric contractions,” European Journal of Applied Physiology and Occupational Physiology, vol. 67, pp. 335–341, Oct. 1993.

[43] M. T. Crow and M. J. Kushmerick, “Chemical energetics of slow- and fast-twitch muscles of the mouse.,” Journal of General Physiology, vol. 79, pp. 147–166, Jan. 1982.

[44] S. M. Baylor and S. Hollingworth, “Intracellular calcium movements during excitation–contraction coupling in mammalian slow-twitch and fast-twitch muscle fibers,” Journal of General Physiology, vol. 139, pp. 261–272, Apr. 2012.

[45] A. V. Terekhov, Y. B. Pesin, X. Niu, M. L. Latash, and V. M. Zatsiorsky, “An analytical approach to the problem of inverse optimization with additive objective functions: an application to human prehension,” Journal of Mathematical Biology, vol. 61, pp. 423–453, Sept. 2010.

[46] D. G. Thelen, F. C. Anderson, and S. L. Delp, “Generating dynamic simulations of movement using computed muscle control,” Journal of biomechanics, vol. 36, no. 3, pp. 321–328, 2003.

[47] A. Annerino, M. Faltas, M. Srinivasan, and P.-I. Gouma, “Towards skin-acetone monitors with selective sensitivity: Dynamics of pani-ca films,” Plos one, vol. 17, no. 4, p. e0267311, 2022.

[48] R. Khusainov, D. Azzi, I. E. Achumba, and S. D. Bersch, “Real-time human ambulation, activity, and physiological monitoring: Taxonomy of issues, techniques, applications, challenges and limitations,” Sensors, vol. 13, no. 10, pp. 12852–12902, 2013.

[49] J. M. Brockway, “Derivation of formulae used to calculate energy expenditure in man,” Human Nutrition. Clinical Nutrition, vol. 41, pp. 463–471, Nov. 1987.

[50] S. Ambike, V. M. Zatsiorsky, and M. L. Latash, “Processes underlying unintentional finger-force changes in the absence of visual feedback,” Experimental Brain Research, vol. 233, pp. 711–721, Mar. 2015.

