## Supplementary Appendix 1 for "Metabolic cost for isometric force scales nonlinearly and predicts how humans distribute forces across limbs"

[*]Co-first authors


### 1 Expressing joint torque in terms of external forces on the limb in the cost function

Consider the single segment model of a limb applying an external force as in Figure **??** and 1. The moment balance about the pivot gives:

$$\tau = \mathrm{L}\ \sin\theta \left(\mathrm{F} - \frac{\mathrm{mg}\ \mathrm{L_{com}}}{\mathrm{L}}\right).$$

Defining $\beta = \frac{\mathrm{mg}\ \mathrm{L_{com}}}{\mathrm{L}}$, we get

$$\tau = \mathrm{L}\,(\mathrm{F} - \beta)\sin\theta. \tag{1.1}$$

We substitute this torque expression in the joint-torque-based energy cost function corresponding to when the left and right limbs together produce the external force. We get:

$$\begin{aligned}\text{Cost function} &= \tau_{\text{left}}^{\gamma} + \tau_{\text{right}}^{\gamma} \\ &= (\mathrm{L}\ \sin\theta)^{\gamma}\left[(\mathrm{F_{left}} - \beta_{\text{left}})^{\gamma} + (\mathrm{F_{right}} - \beta_{\text{right}})^{\gamma}\right],\end{aligned}$$

assuming $\mathrm{L_{left}} = \mathrm{L_{right}} = \mathrm{L}$ and $\theta_{\text{left}} = \theta_{\text{right}} = \theta$. For the purposes of optimisation with a fixed static configuration, the term $(\mathrm{L}\ \sin\theta)^{\gamma}$ does not matter. So, we have:

$$\text{Cost function} = (\mathrm{F_{left}} - \beta_{\text{left}})^{\gamma} + (\mathrm{F_{right}} - \beta_{\text{right}})^{\gamma}. \tag{1}$$

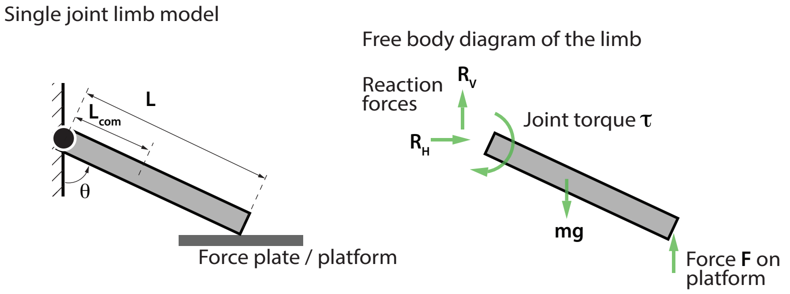


**Figure 1:** Single joint model of a limb used for bilateral force sharing.
